# Development and evaluation of a new methodology for Soft Tissue Artifact compensation in the lower limb

**DOI:** 10.1101/2021.02.08.430265

**Authors:** Bhrigu K. Lahkar, Pierre-Yves Rohan, Ayman Assi, Helene Pillet, Xavier Bonnet, Patricia Thoreux, Wafa Skalli

**Author notes:** Corresponding author: Bhrigu K. Lahkar, Institut de Biomécanique Humaine Georges Charpak, Arts et Métiers ParisTech, 151 Boulevard de l’Hôpital, 75013 Paris, France.

## Abstract

Skin Marker (SM) based motion capture is the most widespread technique used for motion analysis. Yet, the accuracy is often hindered by Soft Tissue Artifact (STA). This is a major issue in clinical gait analysis where kinematic results are used for decision-making. It also has a considerable influence on the results of rigid body and Finite Element (FE) musculoskeletal models that rely on SM-based kinematics to estimate muscle, contact and ligament forces. Current techniques designed to compensate for STA, in particular multi-body optimization methods, assume anatomical simplifications to define joint constraints. These methods, however, cannot adapt to subjects’ bone morphology, particularly for patients with joint lesions, nor easily can account for subject- and location-dependent STA. In this perspective, we propose to develop a conceptual FE based model of the lower limb for STA compensation and evaluate it for 66 healthy subjects under level walking motor task.

Both hip and knee joint kinematics were analyzed, considering both rotational and translational joint motion. Results showed that STA caused underestimation of the hip joint kinematics (up to 2.2°) for all rotational DoF, and overestimation of knee joint kinematics (up to 12°) except in flexion/extension. Joint kinematics, in particular the knee joint, appeared to be sensitive to soft tissue stiffness parameters (rotational and translational mean difference up to 1.5° and 3.4 mm). Analysis of the results using alternative joint representations highlighted the versatility of the proposed modeling approach. This work paves the way for using personalized models to compensate for STA in healthy subjects and different activities.

## 1. Introduction

Accurate assessment of *in vivo* kinematics is essential for providing insights into normal joint functionality (Akbarshahi et al., 2010) and investigation of lower limb joint pathology (Andriacchi and Alexander, 2000). Skin Marker (SM) based motion capture is the most widespread technique used for estimating skeletal kinematics of the lower limb. However, the accuracy of such technique is affected by the relative movement of soft tissues with respect to the underlying bone; a bias commonly referred to as Soft Tissue Artifact (STA). If not compensated for, STA can lead average kinematic errors up to 16 mm in translation and 13° in rotation for the knee joint (Benoit et al., 2006). Such errors may significantly influence the assessment of pathology or the treatment effects in clinical gait analysis (Seffinger and Hruby, 2007).

Different methods have been proposed in the literature to reduce the effect of STA on bone pose estimation (e.g., single-body optimization (Chèze et al., 1995), double anatomical landmark calibration (Cappello et al., 1997), point cluster technique (Andriacchi et al., 1998), and Multi-body Optimisation (MBO) (Lu and O’Connor, 1999)). Amongst these, MBO, which relies on a predefined kinematic model with specific joint constraints, is increasingly used. Initially, simple kinematic constraints such as hinge or spherical joints were considered to represent hip and knee articulation (Charlton et al., 2004; Leardini et al., 2017; Lu and O’Connor, 1999; Reinbolt et al., 2005). Later, anatomical joint constraints (parallel mechanism, coupling curves, ligament length variation, and elastic joint) were introduced, providing encouraging 3D kinematics as they allowed joint displacements (Bergamini et al., 2011; Duprey et al., 2010; Gasparutto et al., 2015; Richard et al., 2016). However, regardless of the joint constraints imposed, generic (unpersonalized) model-derived kinematics were shown inaccurate (knee kinematic error up to 17° and 8 mm) as these models could not adapt to patient-specific geometry, particularly in pathological conditions (Clément et al., 2017). On the other hand, personalization of model geometry based on medical images was shown promising in improving joint kinematics accuracy (Assi et al., 2016; Clément et al., 2015; Nardini et al., 2020).

Joint simplification has indirect consequences on the predictive accuracy of both rigid body musculoskeletal (MSK) models, and Finite Element (FE) based MSK models. Studies that used FE-MSK models to predict local joint mechanics using *in vivo* joint kinematics (Shu et al., 2018; Xu et al., 2016) assumed the knee joint as 1 DoF. Such assumption might result in propagation of uncertainties on the predicted kinematics and would affect the joint reaction as well as muscle and ligament forces.

In light of the aforementioned contexts, reliable estimation of skeletal kinematics with SM-based motion data is still a major challenge (Richard et al., 2017). Furthermore, extensive time and complexity associated with customization of models to subjects’ geometry prohibit large sample size. In that context, methods for 3D reconstruction of bony segments from medical imaging modalities, particularly biplanar X-ray imaging, are promising in research and clinical routine (Chaibi et al., 2012). Also, there is a need for adaptable modeling approaches that can account for subject-, task- and location-dependent STA.

In a previous study, a conceptual FE model was proposed for STA compensation (Skalli et al., 2018). The model consists of bone segments (pelvis, femur and tibia), skin markers, virtual markers, connecting elements between skin markers and corresponding bones, and joint models for the hip and the knee joint. The potential advantage of the proposed model is its versatility with regards to soft tissue stiffness personalization and alternative joint model representation. The objective of the current study was to develop the conceptual model for the lower limb and to implement it on healthy volunteers considering subject-specific models.

## 2. Materials and methods

First, the conceptual model is presented. Then implementation of the model is illustrated within an IRB approved (CEHDF285) study. Finally, the consistency and versatility of the model were investigated through sensitivity of various parameters, including the joints representation.

### 2.1 Conceptual FE model of the lower limb

The rationale underlying the conceptual model is twofold: First, the spring connecting the virtual marker and the skin marker is a simple way of modeling globally and grossly the soft tissue deformation, while being able to adjust the spring stiffness to differentiate both between anatomic regions (such as the pelvis, thigh and shank), and between populations of different skin types (tight or loose). Second, considering virtual markers just beneath the skin markers allows easy post-processing of the results to estimate the corrected position of skin markers. These corrected marker positions are analogous to the model determined markers in standard MBO approaches. Such post-processing helps to use the classical gait analysis software.

The conceptual model of the lower limb consists of bone segments, nodes representing skin markers and virtual markers, joint elements, and elements that connect the skin markers to the corresponding bones. Bone segments are represented by a set of high stiffness (quasi-rigid) beams. The joints between the segments are represented by rigid links, allowing free rotations at the joint and controlled displacements. The connection between a skin marker and the corresponding virtual marker is represented by a linear spring, where all the soft tissue deformation is reported. The connection between the virtual marker and the corresponding bone segment is established with high stiffness beams (Fig. 1).

**Figure 1.**
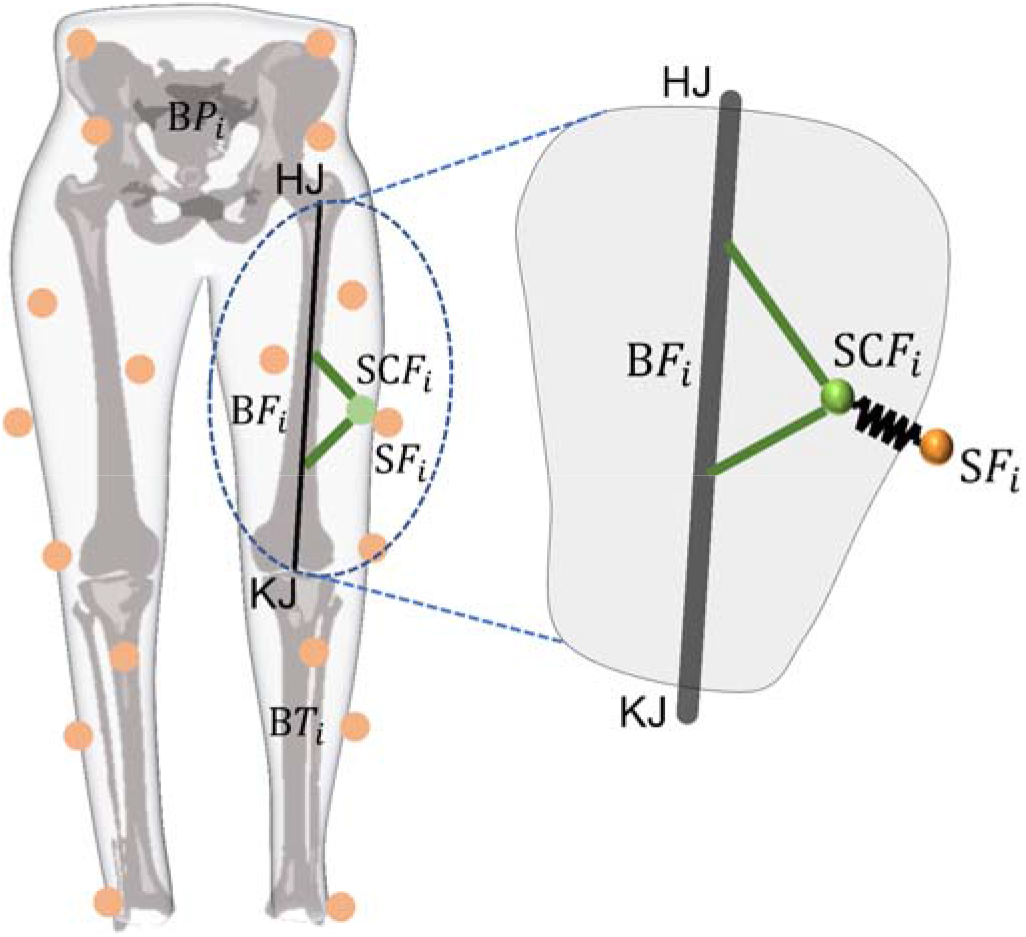
Schematic illustration of the conceptual lower limb FE model. Detailed illustration shown only for the femur segment. B*P*_*i*_, B*F*_*i*_ and B*T*_*i*_ denote pelvis, femur and tibia bone nodes. S*F*_*i*_ and SC*F*_*i*_ are the skin and virtual markers respectively of the for the femur segment. HJ and KJ denote the hip and knee joint respectively.

The proposed model requires only the measured optoelectronic skin marker locations as imposed boundary conditions, without any need for additional force nor any further optimization process. The output of the Finite Element Analysis of the model is the mechanical response resulting in bone motion and the virtual marker positions. From the virtual marker positions, corrected marker positions are obtained.

Skin markers are denoted by *S*, differentiating between those of the pelvis (*SP*), femur (*SF*) and tibia (*ST*). The number of markers for the pelvis (*NMP*), femur (*NMF*) and tibia (*NMT*) is variable and depends on the protocol being considered. Each skin marker is therefore referred to using the corresponding subscript : S*P*_*i*_, S*F*_*i*_ and S*T*_*i*_ respectively for the pelvis (S*P*_1_ to S*P*_*NMP*_), femur (S*F*_1_ to S*F*_*NMF*_) and tibia (S*T*_1_ to S*T*_*NMT*_). Using the same convention, virtual markers are denoted SC (SC*P*_*i*_, SC*F*_*i*_ and SC*T*_*i*_ for the pelvis, femur and tibia respectively), and the bone points are denoted as *B* (differentiating between those of the pelvis (*BP*), femur (*BF*) and tibia (*BT*)). These bone points are different bone landmarks, automatically annotated in the 3D models (Chaibi et al., 2012), which serve as nodes for the FE model. As illustrated further in Fig. 2(a), the pelvis bone was represented by 6 nodes (B*P*_1_ to B*P*_6_), the femur by 7 nodes (B*F*_1_ to B*F*_7_), and the tibia by 6 nodes (B*T*_*1*_ to B*T*_6_)(refer supplementary material for details). Beam elements with elastic modulus (E) of 12 GPa (Choi et al., 1990) were used to connect the nodes for each bone segment. Hip and knee joints are denoted by HJ and KJ respectively.

**Figure 2.**
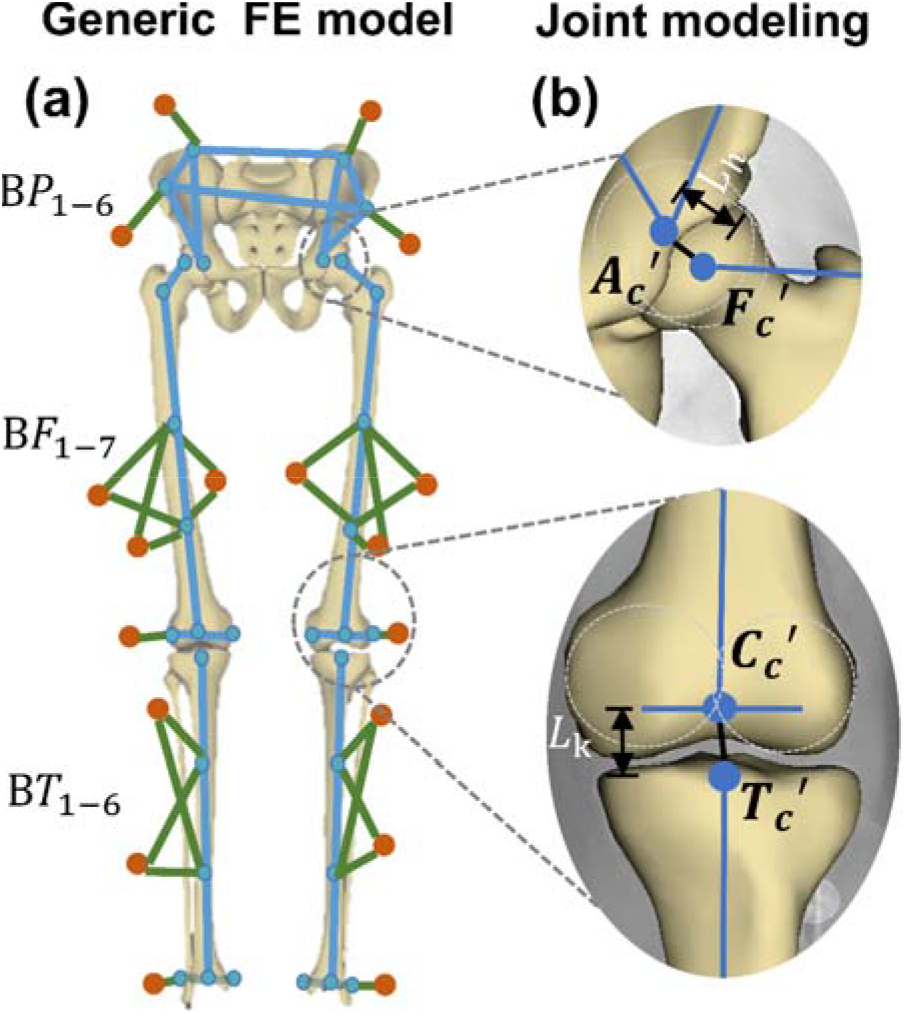
(a) Detailed representation of the lower limb FE model with generic anatomical bony landmarks. Anatomical landmarks for the pelvis (B*P*_l_ to B*P*_6_): right antero-superior iliac spine, right postero-superior iliac spine, left antero-superior iliac spine, left postero-superior iliac spine, and right and left acetabulum centers. For the femur (B*F*_l_ to B*F*_7_): femur head center, greater trochanter, two diaphyseal points, medial and lateral condyle centers and center of the two condyles. For the tibia (B*T*_l_ to B*T*_6_): center of the two plateaus, two diaphyseal points, medial and lateral malleoli and center of the two malleoli. (b) joint modeling of the hip (L_h_) and knee joint (L_k_) with rigid links allowing free rotation and controlled relative displacement.

#### Modeling of the skin marker-bone connection

Each pelvis skin marker (S*P*_1_ to S*P*_4_) was linked to the pelvis bone by a combination of spring element that connects the skin marker to the corresponding virtual marker (SC*P*_1_ to SC*P*_4_), and a beam element that connects the virtual marker to the bone. The springs were assigned with stiffness (k) values in the range 5 kN/m to 65 kN/m (Dumas and Jacquelin, 2017; Gittoes et al., 2006; McLean et al., 2004), whereas the beams were considered highly stiff and assigned the same elastic modulus as that of the bones. The same combination of elements was used to connect the skin markers to the femur and tibia bone segments.

#### Modeling of the joints

As a first option, hip and knee joints were represented each by a rigid link allowing free rotation while controlling the relative displacements (through the length of the link). For the hip joint, the rigid link connected the acetabulum center and femur head center. For the knee joint, the rigid link was defined in the line joining the centroid of the two femoral condyle centers to the centroid of the two tibial plateau centers (Fig. 2(b)) (refer supplementary material for details).

### 2.2. Model implementation

#### 2.2.1. Data Acquisition

66 healthy volunteers were included (age range: 18-60 years; weight: 71.3±15 Kg; height: 170±10 cm) in this study. The only exclusion criterion was previous record of orthopedic surgery of the lower limbs.

Quantitative Movement Analysis was performed on an optoelectronic analysis system comprising 7 video-cameras (Vicon Motion System Ltd., Oxford Metrics, UK). The optoelectronic markers were positioned following the Plug-in Gait^®^ method (Davis et al., 1991), and participants were asked to perform level walking at self-selected speed (Fig. 3(a)). Biplanar radiographs were then acquired using the EOS system (EOS Imaging, France). 3D digital models of bones were obtained using a 3D reconstruction algorithm validated by previous studies (Fig. 3(b)) (Chaibi et al., 2012). The location of skin markers was also computed from biplanar X-Rays.

**Figure 3.**
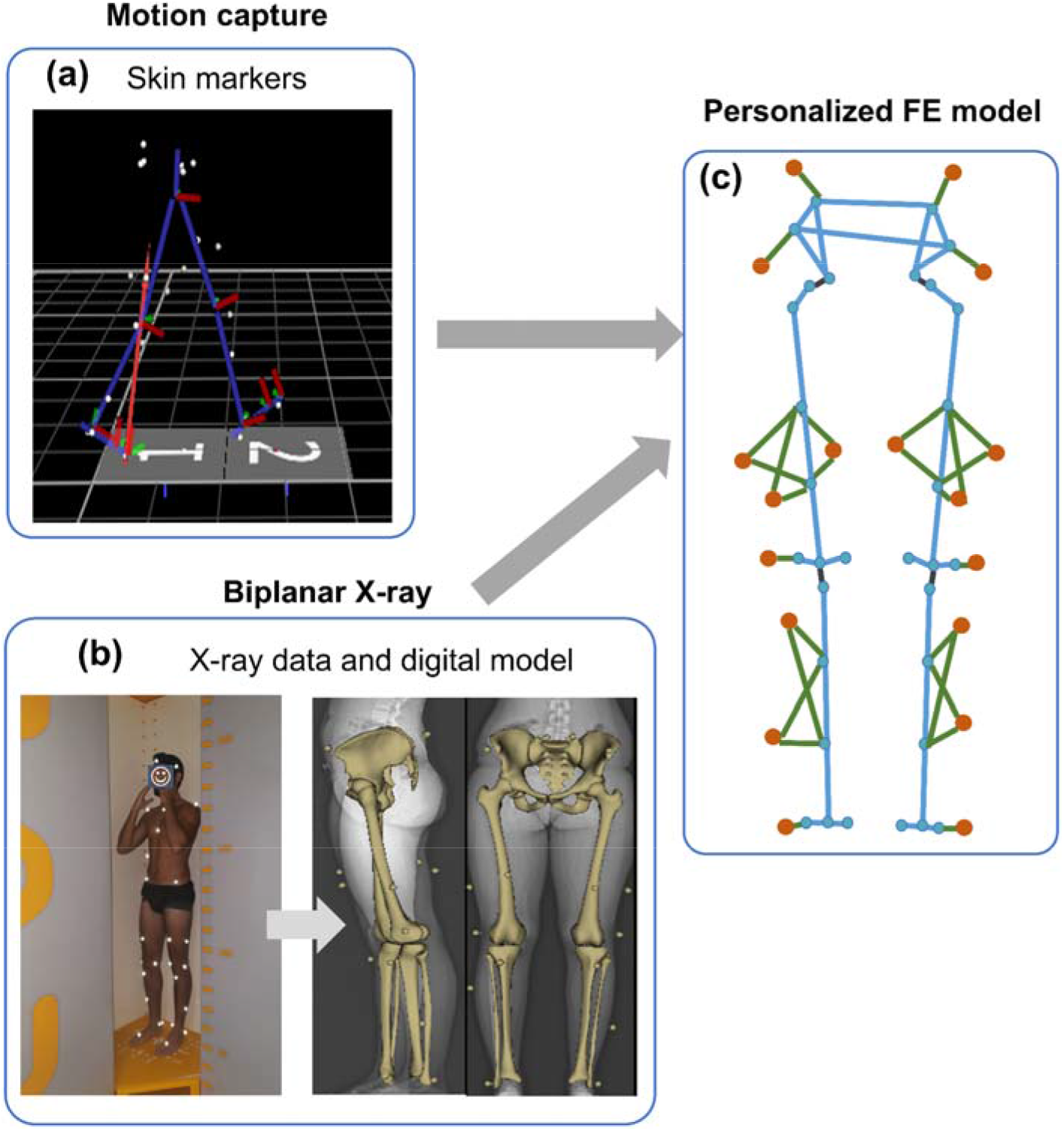
Schematic illustration of FE model personalization (a) the locations of the skin markers throughout gait cycle obtained from motion capture and (b) 3D digital models of the pelvis, femur and tibia built from two orthogonal radiographs (c) anatomical landmarks were identified from the 3D digital models resulting the nodal coordinates of each bone.

#### 2.2.2. Subject-specific FE model development and simulation

From the 3D digital models of the bones, subject-specific anatomical landmarks were automatically identified, resulting in nodal coordinates of each bone, as represented in Fig. 3(c). The distance between the skin and the corresponding virtual marker was arbitrarily chosen as 1 mm (i.e., spring length). A rigid link represented the hip and knee joints. The choice of the rigid link (*L_k_*) for the knee joint was motivated by the *in vitro* kinematic response obtained in a previous study on 12 cadaveric specimens showing that overall knee translations were nearly 20 mm (Rochcongar et al., 2016). For the hip (*L_h_*), the joint length was fixed to 1 mm based on an unpublished data on the hip joint translation quantified using biplanar X-rays. For simplicity, stiffness parameter of the springs was kept constant across all segments and assigned 50 kN/m.

The measured skin marker displacements at each frame from the motion capture were introduced to the model as a prescribed boundary condition. A solution was computed at each frame using commercial FE package ANSYS^®^, with a default Newton-Raphson algorithm, an implicit scheme widely used in numerical procedures for Partial Differential Equations (Bathe, 1996).

#### 2.2.3. Kinematic computation

The positions of the resulting bone segments and virtual markers were used to define corrected markers (*CF*_*i*_) with the same consideration as those of the virtual markers, i.e., rigid links with the bone segment (Fig. 4). The positions of the corrected markers were used to compute STA Compensated (STAC) joint kinematics. Hip and knee joint rotational kinematics were expressed in the pelvis and femur anatomical reference frames (EOS-based) respectively, and with Cardan sequence *YX*’*Z*’’. Hip joint translation was defined as the relative displacement between points *A*_*C*_’ and *F*_*C*_’ expressed in the pelvis anatomical reference frame. Similarly, knee joint translation was defined as a relative displacement between points *C*_*C*_’ and *T*_*C*_’ Iexpressed in the femur anatomical reference frame. Anatomical reference frames were defined as described in (Schlatterer et al., 2009) for the femur and tibia, and in (Dubois, 2014) for the pelvis (Fig. 4). A customized Matlab (MathWorks, Massachusetts, United States) routine was used for both SM-based and STAC kinematic processing. In each case, before and after STA compensation, joint kinematics were obtained using an internal procedure implemented in our previous studies (Azmy et al., 2010; Pillet et al., 2016; Rochcongar et al., 2016). Briefly, skin marker coordinate systems were registered on the bone anatomic reference frames to get the joint kinematics.

**Figure 4.**
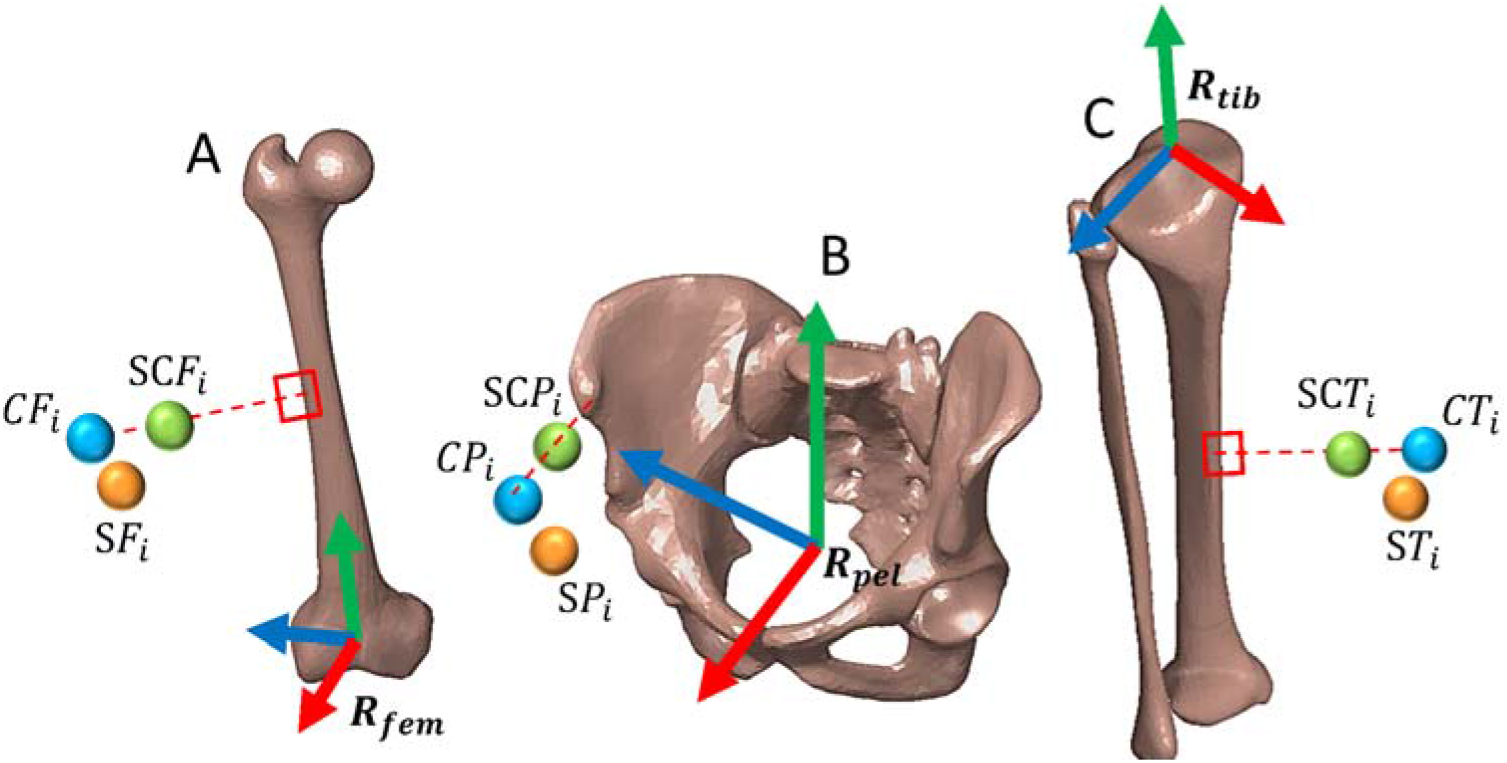
Illustration of corrected markers (*CF*_*i*_, *CP*_*i*_, *CT*_*i*_) and anatomical reference frames (*R*_*fem*_, *R*_*pel*_, *R*_*tib*_) for the (A) femur, (B) pelvis and (C) tibia respectively. Corrected markers are obtained from the virtual marker in a direction orthogonal to the bone segment and 1mm away from virtual marker. Anatomical reference frames for the femur and tibia were defined as described in (Schlatterer et al., 2009) and for pelvis (Dubois, 2014).

Joint kinematics (mean±1SD) for the hip and knee joint were plotted for all DoFs over time normalized gait cycle. The difference in range of motions (dROM) was also computed between SM-based and STAC kinematics.

### 2.3. Illustration of versatility

#### 2.3.1. Sensitivity of spring stiffness and joint length

Two different stiffness values for the springs were implemented (5 kN/m and 65 kN/m) to investigate the influence of stiffness parameters on joint kinematics.

Furthermore, two different knee joint lengths (L_k_ = 21 mm and 31 mm) were arbitrarily considered to investigate the impact of joint lengths on estimated kinematics. Implemented knee joint lengths were based on the minimum and maximum value found in the population. In this case, spring stiffness was kept constant with a value of 65 kN/m.

Differences between the two groups with different spring stiffness values and then joint lengths were analyzed with a Student’s *t*-test or Wilcoxon sign-rank test depending on the outcomes of the Shapiro-Wilk test of normality, using a customized Matlab routine. For all the tests, the significance level was set 0.05 *a priori.*

#### 2.3.2. Alternative joint representations

Two other alternative joint models were considered to illustrate the versatility of the lower limb FE model.

These joint models were:

##### Parallel Mechanism

The centers of the medial and lateral condyles and corresponding tibial plateaus were considered to model the knee joint with two rigid links approximating the femur-tibia contact behavior. The hip joint model was left unaltered (single-link model).

##### Spherical joint

Spherical joint model at the hip and knee joint was considered. The joint constraint location was placed on the femur head center for the hip joint and the mid-point of the two femoral condyles for the knee joint. Such consideration was similar, as reported in previous studies (Sauret et al., 2016).

Differences between alternative joint models were analyzed with a Student’s *t*-test or Wilcoxon sign-rank test depending on the outcomes of the Shapiro-Wilk test of normality, using a customized Matlab routine. For all the tests, the significance level was set 0.05 *a priori.* Spring stiffness value of 65 kN/m was assigned for all the joint models.

### 2.4. Model comparison with multi-body optimization

As no reference kinematics (artifact-free motion) were available, the FE model results were compared to a standard MBO method with spherical joint modeling for both the hip and knee joints (Lu and O’Connor, 1999). The joint constraints and locations incorporated in the MBO were in accordance with the FE model. To compare the kinematic results of the subject-specific FE models with MBO, the same anatomical reference fra mes were defined for the MBO bone segments.

Differences between the two methods were analyzed with a Student’s *t*-test or Wilcoxon sign-rank test depending on the outcomes of the Shapiro-Wilk test of normality, using a customized Matlab routine. For all the tests, the significance level was set 0.05 *a priori.*

## 3. Results

Each FE model with 5 DoF joints required less than 45 sec of run time on a single processor desktop PC to simulate a complete gait cycle of approximately 200–300 frames. All results are synthesized in Table 1.

### 3.1. Joint kinematics

Both rotational and translational kinematics estimated with skin marker measurements and FE model embedding the 5 DoF joint model are illustrated in Figs. 5 and 6 for the hip and knee joints respectively. The joint kinematics are plotted over time-normalized gait cycle.

**Figure 5.**
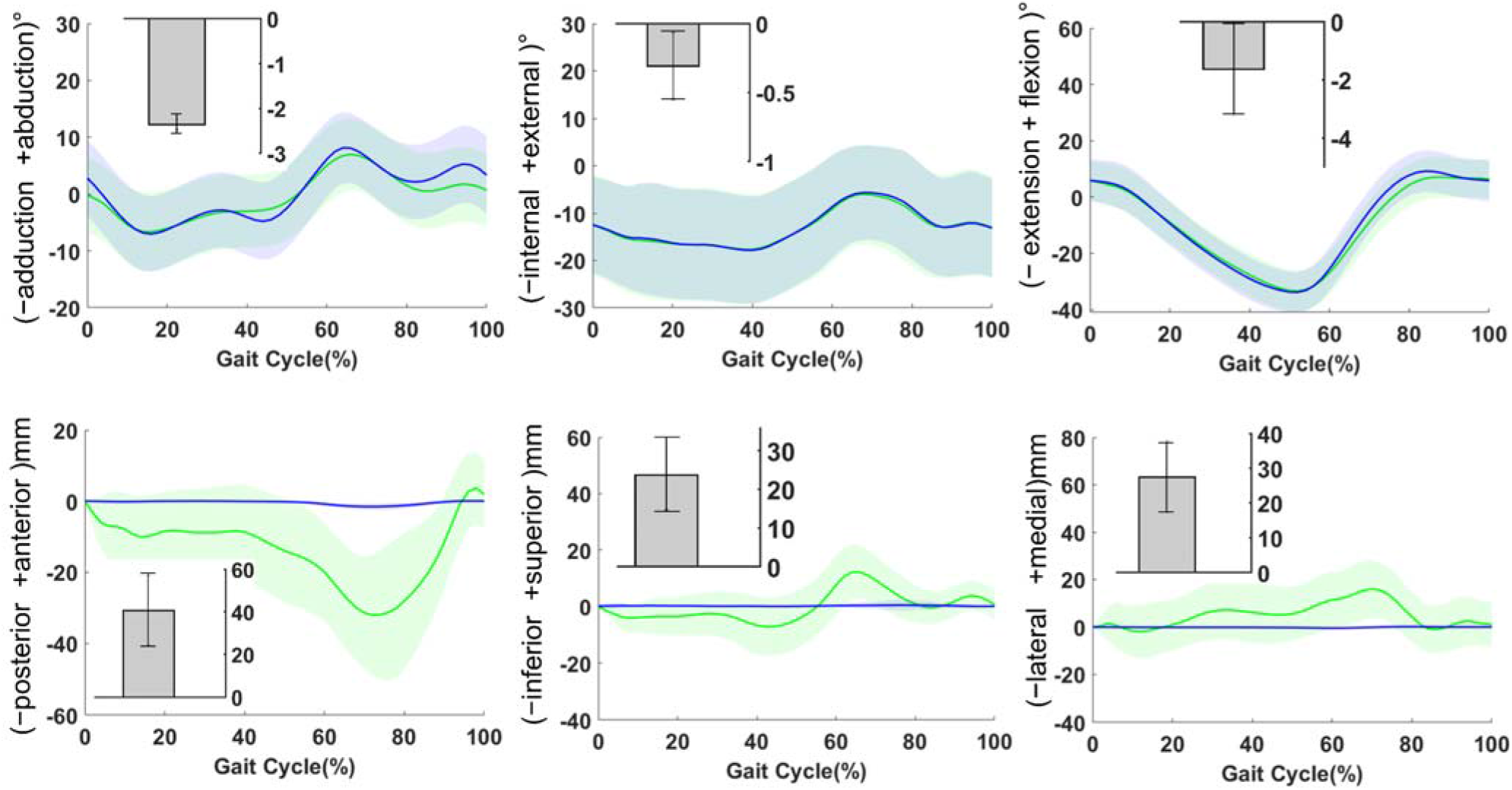
Hip joint kinematics during gait presented as Mean±1SD. Mean values for skin marker-based (green) and FE model predicted results (blue) are shown as solid lines, while standard deviation in lighter shades. Differences in ROM (dROM) between SM-based and STAC results are depicted as insets for all DoFs

**Figure 6.**
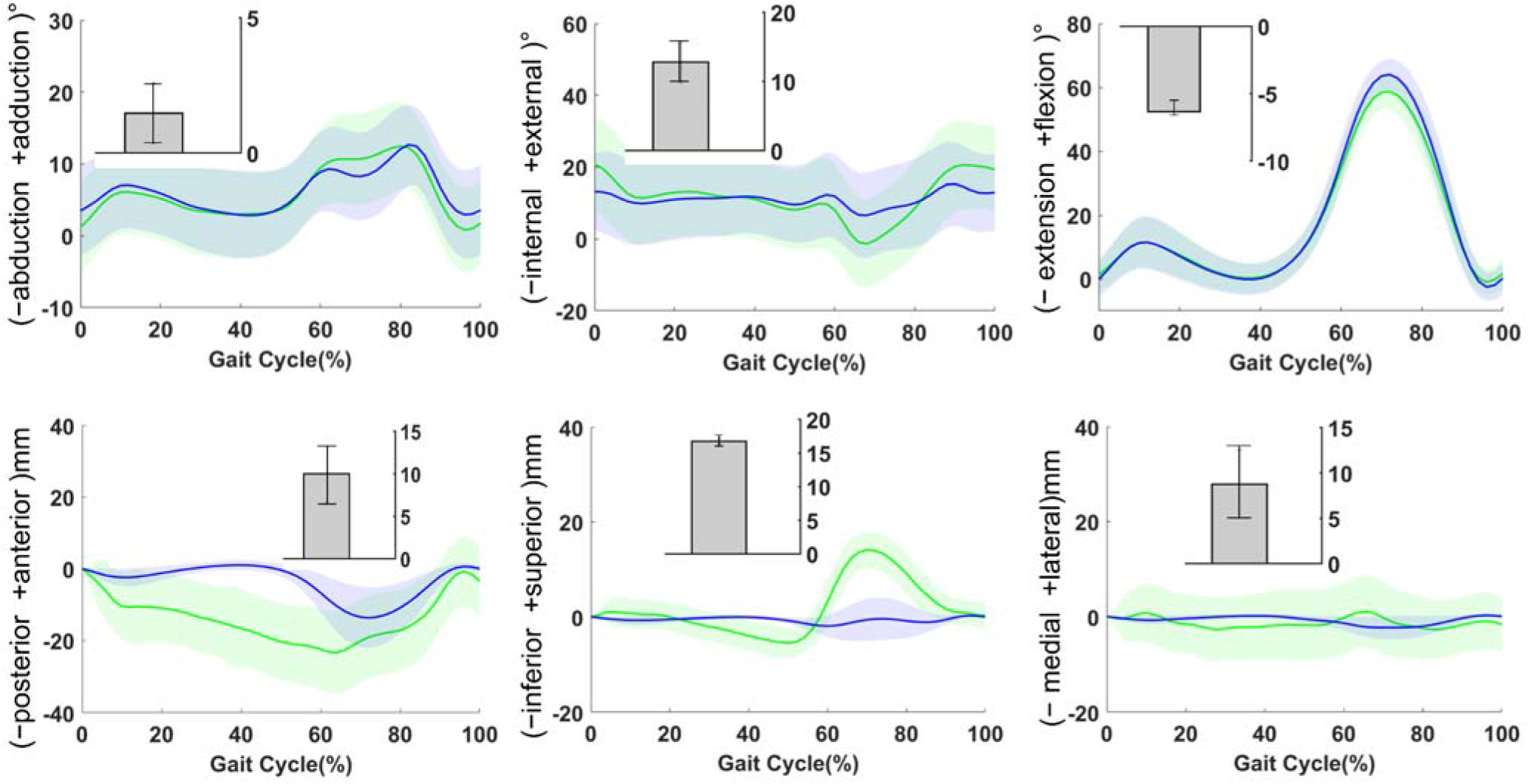
Knee joint kinematics during gait presented as Mean±1SD. Mean values for skin marker-based (green) and FE model predicted results (blue) are shown as solid lines, while standard deviation in lighter shades. Differences in ROM (dROM) between SM-based and STAC results are depicted as insets for all DoFs.

Overall, for the hip joint, STAC and SM-based kinematics exhibited qualitatively similar pattern. However, differences in range of motion (dROM) varied across all DoFs, with a maximum value of −2.2° for Abduction/Adduction (Abd/Add) followed by −1.6° and −0.3° for Flexion/Extension (Flex/Ext) and Internal/External (Int/Ext) rotation respectively. Maximum joint displacement up to 1 mm was observed for STAC kinematics while showing up to 41.5 mm for SM-based kinematics in Posterior/Anterior (Post/Ant) direction. In the Medial/Lateral (Med/Lat) and Inferior/Superior (Inf/Sup) direction, joint displacement exhibited less than 1 mm, whereas SM-based kinematics showed up to 28 mm.

For the knee joint, maximum dROM value was observed for Int/Ext (12.5°) followed by Flex/Ext (−6.3°) and Abd/Add (1.5°) rotation respectively. A maximum of 20 mm of joint displacement was noted in (Post/Ant) direction, while remaining DoFs showed up to 9 mm (Med/Lat) and 3 mm (Inf/Sup) for STAC kinematics. These results showed up to 30.5 mm, 12.5 mm, and 21 mm respectively, for SM-based kinematics.

### 3.2. Sensitivity study

#### 3.2.1. Spring stiffness parameter and joint length

With two different values of spring stiffness parameters (5 kN/m and 65 kN/m), no statistical significance in dROM was noted for the hip joint kinematics. As for the knee joint, different spring stiffness revealed significant dROM for all DoFs except Flex/Ext.

With different knee joint lengths (21 mm and 31 mm), hip translational kinematics displayed significant variability for Lat/Med motions (by 2 mm), while showing less than 1 mm change for remaining DoFs. As for knee translational kinematics, significant dROM was observed for the Post/Ant and Inf/Sup motions.

#### 3.2.2. Influence of alternative joint models on kinematics

Different joint representations displayed varying kinematic changes across all DoFs for the hip and knee joints. Among hip joint kinematic results, significant dROM was observed only for the Lat/Med (up to 4 mm) and Post/Ant motion (up to 2.7 mm). Similarly, knee joint kinematics significantly varied up to 3.4° in dROM for Int/Ext rotation when joint model was altered, along with Post/Ant motion (up to 7.6 mm), for the single link Vs parallel mechanism.

### 3.3. FE Model comparison with MBO

Statistically significant differences between MBO-based and FE-based STA compensation for the hip and knee joints were found (p<0.05). Those differences, however, were always in the range of 0.7° to 2°.

## 3. Discussion

Soft Tissue Artifact compensation is essential for accurate estimation of *in vivo* joint kinematics in both research and clinical routine; however, personalization and versatility of current model-based methods still represent a challenge. The purpose of this study was to develop and evaluate a conceptual FE model of the lower limb for STA compensation. The proposed method was evaluated on a population of 66 subjects. This model is computationally fast (less than 45 sec run time), and its main advantage is versatility allowing a wide range of parameters and joint representations to be considered.

Qualitatively similar kinematic patterns were observed between SM-based and FE-based STA compensated (STAC) results for both the hip and knee joints, with differences in range ROM across all DoFs. SM-based kinematics were comparable to the literature (D’Isidoro et al., 2020; Fiorentino et al., 2017). Results obtained showed that overall rotational ROM was underestimated by SM-based results up to 2.2° for the hip joint, thus confirming similar observations reported in studies that compared SM-based ROM to dual fluoroscopic measurements (Fiorentino et al., 2020). For the knee joint, SM-based ROM was smaller by 6.3° for the Flex/Ext, whereas other DoFs revealed up to 12° higher as compared to STAC kinematics. For translational kinematics of the knee joint, SM-based results were higher as compared to STAC kinematics. STAC knee kinematics were comparable to studies that reported either bone-pin-based or fluoroscopy-based kinematics (Benoit et al., 2006; Kozanek et al., 2009; Myers et al., 2012). Nevertheless, we observed overall higher dROM values between SM-based and STAC as compared to the studies that reported in the range 4.4°–5.3° for rotational kinematics, and up to 13 mm for translational kinematics (Benoit et al., 2006; Leardini et al., 2005). These discrepancies may arise from the experimental protocol, such as number of markers, cluster configuration and location.

Sensitivity study showed that joint kinematics (particularly the knee joint) were sensitive to spring stiffness exhibiting dROM value up to 1.5° for the rotational kinematics and up to 3.5 mm for translational kinematics. Different joint representations revealed that alternative joint models have considerable influence on the estimated kinematics, particularly knee Int/Ext rotation (up to 3.4°) and translations (up to 7.6 mm), establishing similar remarks as reported in the literature (Duprey et al., 2010; Richard et al., 2017).

Joint kinematics computed using the proposed FE-based STA compensation model were able to consider the joint simplifications used in a standard MBO method in the literature, producing consistent results. However, the proposed approach aims to overcome the limitations of the MBO method underlined by (Clément et al., 2017), who showed that simplified joint constraints (kinematic or anatomical) were inadequate for clinical applications. Indeed, such simplifications considered in MBO methods may be sufficient for many movement analysis applications; however, this is not the case for pathological cases. For example, in a previously unpublished study using EOS imaging, the translations at the hip joint (distance between the femur head and acetabulum center) during a change of posture from standing to sitting was quantified varying up to 5 mm in pathological population. In such cases, it is apparent that standard joint representation, such as a spherical joint, is not relevant. The main advantage of the new procedure is its versatility. Indeed, it can be modified to incorporate more or less detailed joint representations to improve joint mechanics estimation. It could also evolve into subject-specific modeling for clinical applications. For example, the spring stiffness could be personalized based on quantitative soft tissue deformation that can be assessed using medical images (Südhoff et al., 2007). Such avenues are currently under investigation.

This study has some limitations. First, there was no reference kinematics to compare the results to. Therefore, the joint kinematics exhibited by the FE models were compared to those computed with the MBO method. Nevertheless, as we cannot consider MBO as a fully reliable solution for STA compensation (Richard et al., 2017), such comparison is only for assessing the qualitative performance of the FE model. Second, STA parameters implemented in the model were arbitrary as there is a lack of data in the literature. Personalization of such parameters is, however, essential to encompass different range of subjects (young, adult, patients with CP and OA etc.). Third, joint representation in this model is still simplified, which could be insufficient for investigating local joint mechanics for healthy or pathological joints (Adouni et al., 2012; Lenhart et al., 2015; Shu et al., 2018; Valente et al., 2014). Nevertheless, as a preliminary step, the current contribution only focused on exploring and facilitating personalization of the parameters that are important for STA compensation. Moreover, even with simplified joint representation, the model could limit the effect of STA in joint kinematics. Fourth, although the proposed approach may give the impression that it complicates the process of STA compensation in gait analysis, the perspectives are numerous, as already highlighted. Fifth, no external forces nor inertial/mass forces were imposed on the model. The only boundary conditions were the external skin markers displacements. Considering the inertial/mass forces would be necessary when dynamic phenomena are essential to take into account, for example, in sports biomechanics. Finally, the study was based on a single motor task, i.e., level walking. Therefore, the results may vary with other motor tasks (hopping, cutting, stand-to-sit) and hence the interpretations.

The proposed approach may serve in two major fields of applications: First, in gait analysis for research, where classical scaling techniques are used to obtain subject-specific geometry instead of image-based model personalization. Compensating for STA with such method is possible with an approximated model geometry, while being able to differentiate soft tissue stiffness parameters between different sub-groups. To be noted that as such scaling techniques often consider gross anthropometry of the subject and disregards distinctive features of the joint, therefore can be considered “not actually personalized” (Nardini et al., 2020; Smale et al., 2019). Second field of application can be clinical gait analysis, where image-based model personalization could capture anatomical details of the joint structures.

In conclusion, as a first study, we presented a conceptual FE model of the lower limb for STA compensation and evaluated it in a population of 66 subjects with varying morphologies. The model appeared to be satisfactory in compensating for STA and versatile, facilitating parameters necessary for model personalization. The methodology developed and evaluated in this study may improve the accuracy of kinematic predictions, which is instrumental for MSK models as well as making clinical decisions. In the current contribution, the human model used for the computations consists only of the lower limbs (pelvis, femur and tibia). However, the same approach can be considered for the whole body, which could be particularly interesting for the shoulder joint (Duprey et al., 2017). Future work could focus on further model evaluation based on *in vivo* data, such as dual fluoroscopy.

## Conflict of Interest

None

## Acknowledgments

The authors are deeply grateful to the ParisTech BiomecAM chair program on subject-specific musculoskeletal modeling for financial support.

## Notes

### Competing Interest Statement

The authors have declared no competing interest.

## References

Adouni, M., Shirazi-Adl, A., Shirazi, R., 2012. Computational biodynamics of human knee joint in gait: From muscle forces to cartilage stresses. J Biomech 45, 2149–2156. https://doi.org/10.1016/j.jbiomech.2012.05.040

Akbarshahi, M., Schache, A.G., Fernandez, J.W., Baker, R., Banks, S., Pandy, M.G., 2010. Non-invasive assessment of soft-tissue artifact and its effect on knee joint kinematics during functional activity. J Biomech 43, 1292–1301. https://doi.org/10.1016/j.jbiomech.2010.01.002

Andriacchi, T.P., Alexander, E.J., 2000. Studies of human locomotion: Past, present and future. J Biomech 33, 1217–1224. https://doi.org/10.1016/S0021-9290(00)00061-0

Andriacchi, T.P., Alexander, E.J., Toney, M.K., Dyrby, C., Sum, J., 1998. A Point Cluster Method for In Vivo Motion Analysis: Applied to a Study of Knee Kinematics. J Biomech Eng 120, 743–749. https://doi.org/10.1115/1.2834888

Assi, A., Sauret, C., Massaad, A., Bakouny, Z., Pillet, H., Skalli, W., Ghanem, I., 2016. Validation of hip joint center localization methods during gait analysis using 3D EOS imaging in typically developing and cerebral palsy children. Gait Posture 48, 30–35. https://doi.org/10.1016/j.gaitpost.2016.04.028

Azmy, C., Guérard, S., Bonnet, X., Gabrielli, F., Skalli, W., 2010. EOS^®^ orthopaedic imaging system to study patellofemoral kinematics: Assessment of uncertainty. Orthop Traumatol Surg Res 96, 28–36. https://doi.org/10.1016/j.otsr.2009.10.013

Bathe, K.J., 1996. Finite Element Procedures, Englewood Cliffs New Jersey.

Benoit, D.L., Ramsey, D.K., Lamontagne, M., Xu, L., Wretenberg, P., Renström, P., 2006. Effect of skin movement artifact on knee kinematics during gait and cutting motions measured in vivo. Gait Posture 24, 152–164. https://doi.org/10.1016/j.gaitpost.2005.04.012

Bergamini, E., Pillet, H., Hausselle, J., Thoreux, P., Guerard, S., Camomilla, V., Cappozzo, A., Skalli, W., 2011. Tibio-femoral joint constraints for bone pose estimation during movement using multi-body optimization. Gait Posture 33, 706–711. https://doi.org/10.1016/j.gaitpost.2011.03.006

Cappello, A., Cappozzo, A., La Palombara, P.F., Lucchetti, L., Leardini, A., 1997. Multiple anatomical landmark calibration for optimal bone pose estimation. Hum Mov Sci 16, 259–274. https://doi.org/https://doi.org/10.1016/S0167-9457(96)00055-3

Chaibi, Y., Cresson, T., Aubert, B., Hausselle, J., Neyret, P., Hauger, O., de Guise, J.A., Skalli, W., 2012. Fast 3D reconstruction of the lower limb using a parametric model and statistical inferences and clinical measurements calculation from biplanar X-rays. Comput Methods Biomech Biomed Engin 15, 457–466. https://doi.org/10.1080/10255842.2010.540758

Charlton, I.W., Tate, P., Smyth, P., Roren, L., 2004. Repeatability of an optimised lower body model. Gait Posture 20, 213–221. https://doi.org/10.1016/j.gaitpost.2003.09.004

Chèze, L., Fregly, B.J., Dimnet, J., Sztankóová, Z., Kyselová, J., Rychtářová, J., Czerneková, V., 1995. A solidification procedure to facilitate kinematic analyses based on video system data. J Biomech 28, 879–884. https://doi.org/https://doi.org/10.1016/0021-9290(95)95278-D

Clément, J., Dumas, R., Hagemeister, N., de Guise, J.A., 2017. Can generic knee joint models improve the measurement of osteoarthritic knee kinematics during squatting activity? Comput Methods Biomech Biomed Engin 20, 94–103. https://doi.org/10.1080/10255842.2016.1202935

Clément, J., Dumas, R., Hagemeister, N., de Guise, J.A., 2015. Soft tissue artifact compensation in knee kinematics by multi-body optimization: Performance of subject-specific knee joint models. J Biomech 48, 3796–3802. https://doi.org/10.1016/j.jbiomech.2015.09.040

D’Isidoro, F., Brockmann, C., Ferguson, S.J., 2020. Effects of the soft tissue artefact on the hip joint kinematics during unrestricted activities of daily living. J Biomech 109717. https://doi.org/10.1016/j.jbiomech.2020.109717

Davis, R.B., Õunpuu, S., Tyburski, D., 1991. A gait analysis data collection and reduction technique. Hum Mov Sci 10, 575–587. https://doi.org/https://doi.org/10.1016/0167-9457(91)90046-Z

Dubois, G., 2014. Contribution à la modélisation musculo-squelettique personnalisée du membre inférieur combinant stéréoradiographie et ultrason.

Dumas, R., Jacquelin, E., 2017. Stiffness of a wobbling mass models analysed by a smooth orthogonal decomposition of the skin movement relative to the underlying bone. J Biomech 62, 47–52. https://doi.org/10.1016/j.jbiomech.2017.06.002

Duprey, S., Cheze, L., Dumas, R., 2010. Influence of joint constraints on lower limb kinematics estimation from skin markers using global optimization. J Biomech 43, 2858–2862. https://doi.org/10.1016/j.jbiomech.2010.06.010

Duprey, S., Naaim, A., Moissenet, F., Begon, M., Chèze, L., 2017. Kinematic models of the upper limb joints for multibody kinematics optimisation: An overview. J Biomech 62, 87–94. https://doi.org/10.1016/j.jbiomech.2016.12.005

Fiorentino, N.M., Atkins, P.R., Kutschke, M.J., Bo Foreman, K., Anderson, A.E., 2020. Soft Tissue Artifact Causes Underestimation of Hip Joint Kinematics and Kinetics in a Rigid-Body Musculoskeletal Model. J Biomech 108, 109890. https://doi.org/10.1016/j.jbiomech.2020.109890

Fiorentino, N.M., Atkins, P.R., Kutschke, M.J., Goebel, J.M., Foreman, K.B., Anderson, A.E., 2017. Soft tissue artifact causes significant errors in the calculation of joint angles and range of motion at the hip. Gait Posture 55, 184–190. https://doi.org/10.1016/j.gaitpost.2017.03.033

Gasparutto, X., Sancisi, N., Jacquelin, E., Parenti-Castelli, V., Dumas, R., 2015. Validation of a multi-body optimization with knee kinematic models including ligament constraints. J Biomech 48, 1141–1146. https://doi.org/10.1016/j.jbiomech.2015.01.010

Gittoes, M.J.R., Brewin, M.A., Kerwin, D.G., 2006. Soft tissue contributions to impact forces simulated using a four-segment wobbling mass model of forefoot – heel landings 25, 775–787. https://doi.org/10.1016/j.humov.2006.04.003

Kozanek, M., Hosseini, A., Liu, F., Van de Velde, S.K., Gill, T.J., Rubash, H.E., Li, G., 2009. Tibiofemoral kinematics and condylar motion during the stance phase of gait. J Biomech 42, 1877–1884. https://doi.org/10.1016/j.jbiomech.2009.05.003

Leardini, A., Belvedere, C., Nardini, F., Sancisi, N., Conconi, M., Parenti-Castelli, V., 2017. Kinematic models of lower limb joints for musculo-skeletal modelling and optimization in gait analysis. J Biomech 62, 77–86. https://doi.org/10.1016/j.jbiomech.2017.04.029

Leardini, A., Chiari, A., Della Croce, U., Cappozzo, A., 2005. Human movement analysis using stereophotogrammetry Part 3. Soft tissue artifact assessment and compensation. Gait Posture 21, 212–225. https://doi.org/10.1016/j.gaitpost.2004.05.002

Lenhart, R.L., Kaiser, J., Smith, C.R., Thelen, D.G., 2015. Prediction and Validation of Load-Dependent Behavior of the Tibiofemoral and Patellofemoral Joints During Movement. Ann Biomed Eng 43, 2675–2685. https://doi.org/10.1007/s10439-015-1326-3

Lu, T.W., O’Connor, J.J., 1999. Bone position estimation from skin marker co-ordinates using global optimisation with joint constraints. J Biomech 32, 129–134. https://doi.org/10.1016/S0021-9290(98)00158-4

McLean, S.G., Su, A., van den Bogert, A.J., 2004. Development and Validation of a 3-D Model to Predict Knee Joint Loading During Dynamic Movement. J Biomech Eng 125, 864–874. https://doi.org/10.1115/1.1634282

Myers, C.A., Torry, M.R., Shelburne, K.B., Giphart, J.E., Laprade, R.F., Woo, S.L.Y., Steadman, J.R., 2012. In vivo tibiofemoral kinematics during 4 functional tasks of increasing demand using biplane fluoroscopy. Am J Sports Med 40, 170–178. https://doi.org/10.1177/0363546511423746

Nardini, F., Belvedere, C., Sancisi, N., Conconi, M., Leardini, A., Durante, S., Parenti-Castelli, V., 2020. An anatomical-based subject-specific model of in-vivo knee joint 3D kinematics from medical imaging. Appl Sci 10, 8–12. https://doi.org/10.3390/app10062100

Pillet, H., Bergamini, E., Rochcongar, G., Camomilla, V., Thoreux, P., Rouch, P., Cappozzo, A., Skalli, W., 2016. Femur, tibia and fibula bone templates to estimate subject-specific knee ligament attachment site locations. J Biomech 49, 3523–3528. https://doi.org/10.1016/j.jbiomech.2016.09.027

Reinbolt, J.A., Schutte, J.F., Fregly, B.J., Koh, B. Il, Haftka, R.T., George, A.D., Mitchell, K.H., 2005. Determination of patient-specific multi-joint kinematic models through two-level optimization. J Biomech 38, 621–626. https://doi.org/10.1016/j.jbiomech.2004.03.031

Richard, V., Cappozzo, A., Dumas, R., 2017. Comparative assessment of knee joint models used in multi-body kinematics optimisation for soft tissue artefact compensation. J Biomech 62, 95–101. https://doi.org/10.1016/j.jbiomech.2017.01.030

Richard, V., Lamberto, G., Lu, T.W., Cappozzo, A., Dumas, R., 2016. Knee Kinematics Estimation Using Multi-Body Optimisation Embedding a Knee Joint Stiffness Matrix: A Feasibility Study. PLoS One 11, 1–18. https://doi.org/10.1371/journal.pone.0157010

Rochcongar, G., Pillet, H., Bergamini, E., Moreau, S., Thoreux, P., Skalli, W., Rouch, P., 2016. A new method for the evaluation of the end-to-end distance of the knee ligaments and popliteal complex during passive knee flexion. Knee 23, 420–425. https://doi.org/10.1016/j.knee.2016.02.003

Sauret, C., Pillet, H., Skalli, W., Sangeux, M., 2016. On the use of knee functional calibration to determine the medio-lateral axis of the femur in gait analysis: Comparison with EOS biplanar radiographs as reference. Gait Posture 50, 180–184. https://doi.org/10.1016/j.gaitpost.2016.09.008

Schlatterer, B., Suedhoff, I., Bonnet, X., Catonne, Y., Maestro, M., Skalli, W., 2009. Skeletal landmarks for TKR implantations: Evaluation of their accuracy using EOS imaging acquisition system. Orthop Traumatol Surg Res 95, 2–11. https://doi.org/10.1016/j.otsr.2008.05.001

Seffinger, M.A., Hruby, R.J., 2007. CHAPTER 3 - Manual Diagnostic Procedures Overview, in: Seffinger, M.A., Hruby, R.J.B.T.-E.-B.M.M. (Eds.),. W.B. Saunders, Philadelphia, pp. 35–58. https://doi.org/https://doi.org/10.1016/B978-1-4160-2384-5.50007-9

Shu, L., Yamamoto, K., Yao, J., Saraswat, P., Liu, Y., Mitsuishi, M., Sugita, N., 2018. A subject-specific finite element musculoskeletal framework for mechanics analysis of a total knee replacement. J Biomech 77, 146–154. https://doi.org/10.1016/j.jbiomech.2018.07.008

Skalli, W., Hermal, T., Bonnet, X., Assi, A., Pillet, H., 2018. A subject-specific finite element based method for soft tissue artefact reduction in motion analysis. In: 8th World Congress of Biomechanics. URL: https://app.oxfordabstracts.com/events/123/program-app/submission/24056

Smale, K.B., Conconi, M., Sancisi, N., Krogsgaard, M., Alkjaer, T., Parenti-Castelli, V., Benoit, D.L., 2019. Effect of implementing magnetic resonance imaging for patient-specific OpenSim models on lower-body kinematics and knee ligament lengths. J Biomech 83, 9–15. https://doi.org/10.1016/j.jbiomech.2018.11.016

Südhoff, I., Van Driessche, S., Laporte, S., de Guise, J.A., Skalli, W., 2007. Comparing three attachment systems used to determine knee kinematics during gait. Gait Posture 25, 533–543. https://doi.org/10.1016/j.gaitpost.2006.06.002

Valente, G., Pitto, L., Testi, D., Seth, A., Delp, S.L., Stagni, R., Viceconti, M., Taddei, F., 2014. Are subject-specific musculoskeletal models robust to the uncertainties in parameter identification? PLoS One 9. https://doi.org/10.1371/journal.pone.0112625

Xu, C., Silder, A., Zhang, J., Hughes, J., Unnikrishnan, G., Reifman, J., Rakesh, V., 2016. An Integrated Musculoskeletal-Finite-Element Model to Evaluate Effects of Load Carriage on the Tibia during Walking. J Biomech Eng 138, 101001. https://doi.org/10.1115/1.4034216

